# The CYK-4 GAP domain regulates cortical targeting of centralspindlin to promote contractile ring assembly and facilitate ring dissolution

**DOI:** 10.1101/2024.10.29.620943

**Authors:** Aleesa J Schlientz, Kian-Yong Lee, J. Sebastián Gómez-Cavazos, Pablo Lara-González, Arshad Desai, Karen Oegema

## Abstract

During cytokinesis, an equatorial contractile ring partitions the cell contents. Contractile ring assembly requires an equatorial zone of active GTP-bound RhoA generated by the guanine nucleotide exchange factor ECT2^1,2^. ECT2 is activated by centralspindlin, a complex composed of two molecules each of kinesin-6 and CYK4. During anaphase, Centralspindlin is activated at the central spindle between the separating chromosomes and diffuses to the plasma membrane, where it engages with ECT2 via its N-terminal half^3,4^. The C-terminal half of CYK4 contains a lipid-binding C1 domain that contributes to plasma membrane targeting^5^ and a GTPase-activating protein (GAP) domain that has an interaction surface for a Rho family GTPase, whose functions have remained unclear ^1,3,4,6,7^. Here, using the one-cell stage *C. elegans* embryo as a model, we show that RhoA and the Rho-binding interface of the CYK4 GAP domain drive the recruitment of centralspindlin to the equatorial cortex. By contrast, a point mutant that selectively disrupts GAP activity does not prevent cortical centralspindlin recruitment but instead substantially delays dissipation of centralspindlin from the cortex. These findings suggest that positive feedback, in which centralspindlin recruitment promotes the generation of active RhoA and active RhoA drives centralspindlin recruitment, is central to the rapid assembly of the contractile ring within a narrow time window. They also indicate that the CYK4 GAP catalytic activity contributes to release of centralspindlin from the cortex, potentially to ensure timely dissolution of the contractile ring.

## RESULTS AND DISCUSSION

### An approach for robust depletion of CYK-4 in one-cell stage *C. elegans* embryos

A key advantage of the *C. elegans* embryo is the ability to develop molecular replacement systems in which an endogenous protein is depleted by RNA interference (RNAi) and replaced with a wild-type (WT) or mutant version expressed from a single copy transgene under control of its endogenous regulatory elements^8^. This approach has been used to analyze the role of the centralspindlin component CYK-4^9,10^. While depletion sufficient to prevent successful cytokinesis is observed following targeting of *cyk-4* via RNAi^9–11^, precise molecular replacement of CYK-4 is made more challenging by CYK-4’s role in the cytokinesis-like process that buds oocytes from the syncytial germline (**Fig. S1A**)^12^, as *cyk-4(RNAi)* worms become sterile before embryos that are penetrantly depleted of CYK-4 protein are generated. To overcome this challenge and achieve more potent CYK-4 depletion, we developed a 2-step RNAi strategy in strains with a single-copy integrated RNAi-resistant transgene encoding CYK-4::GFP fused to the N-terminus of MEI-1 (aa1-224; CYK-4::GFP::MEI-1N), which contains a degron that promotes protein destruction at the meiosis-to-mitosis transition in the embryo^13,14^ (**Fig. S1B,C**). L4 hermaphrodites were injected 42-48 hours before dissection with dsRNA targeting endogenous *cyk-4*, and CYK-4::GFP::MEI-1N encoded by the RNAi-resistant transgene supported germline function as endogenous CYK-4 was depleted (**Fig. S1D,E**). In the resulting embryos, the MEI-1N tag causes degradation during the transition from meiosis to the first mitotic division; CYK-4::GFP::MEI-1N was further reduced by re-injecting the worms with a dsRNA targeting the *gfp* sequence in the CYK-4::GFP::MEI-1N transgene along with a dsRNA targeting endogenous *cyk-4* during the final 18-24 hours before dissection (*2-step* RNAi; **Fig. S1F**). This 2-step strategy enabled sufficiently robust depletion of CYK-4 to nearly completely eliminate CYK-4 from the central spindle in one-cell stage embryos (**Fig. S1F,G**) and facilitated a thorough evaluation of CYK-4 GAP domain function in cytokinesis.

### Contrasting effects of mutating the Rho GTPase-binding interface versus catalytic GAP activity of CYK-4

The CYK-4 GAP domain is predicted to interact with a Rho family small GTPase. The human CYK4 GAP domain has been crystallized bound to RhoA^15^, and AlphaFold2^16^ predicts a similar interaction between *C. elegans* RhoA (encoded by *rho-1*) and the CYK-4 GAP domain (**Fig. 1A**). To investigate the role of the Rho-binding interface and catalytic activity of the CYK-4 GAP domain, we compared the effects of two GAP domain mutants: a three residue mutation in the Rho-binding interface (R459A, K495A, R499E; GAP RhoInt^mut^), predicted to prevent interaction with Rho GTPases, and an arginine finger mutation (R459A; GAP Cat^mut^), predicted to selectively prevent the ability of the GAP domain to promote GTP hydrolysis. Transgenes encoding WT CYK-4 and a version of CYK-4 with its lipid-binding C1 domain deleted (ΔC1), which is expected to disrupt recruitment of CYK-4 to the cortex^5,10^, were used as controls (**Fig. 1B**). These single-copy untagged RNAi-resistant CYK-4 transgenes expressed under control of the endogenous *cyk-4* regulatory sequences (WT, ΔC1, GAP RhoInt^mut^, and GAP Cat^mut^) were previously functionally characterized in the germline and shown to be expressed at equivalent levels after depletion of endogenous CYK-4^12^. We combined these transgenes with the 2-step approach to generate embryos depleted of CYK-4, as described above. Analysis of chromosome separation after depletion of endogenous CYK-4 confirmed that all of the transgene-encoded CYK-4 variants supported the formation of a central spindle that prevented abrupt separation of chromosomes after anaphase onset (**Fig. S2**). Cytokinesis in the presence of the transgene-encoded CYK-4 variants was characterized by imaging *in situ*-tagged non-muscle myosin II fused to RNAi-resistant GFP (NMY-2::GFP; **Fig. 1C,D**; **Fig. S3A**) or by imaging mCherry-fused probes for the plasma membrane and DNA (**Fig. S3B-C**). As expected, WT CYK-4 supported rapid and complete cytokinetic furrow closure, while the no transgene and ΔC1 controls exhibited 100% early cytokinesis failure in which the furrow never fully closed (**Fig. 1C,D; Fig. S3B,C**). Note that the furrowing that remains when CYK-4 is depleted (**Fig. 1C**, **Fig. S3B**; No transgene) is due to the presence of NOP-1, a *C. elegans*-specific protein that functions in parallel to centralspindlin to promote RhoA activation^17^. While NOP-1 augments RhoA activation and contractility, it is not essential for cytokinesis.

**Figure 1.**
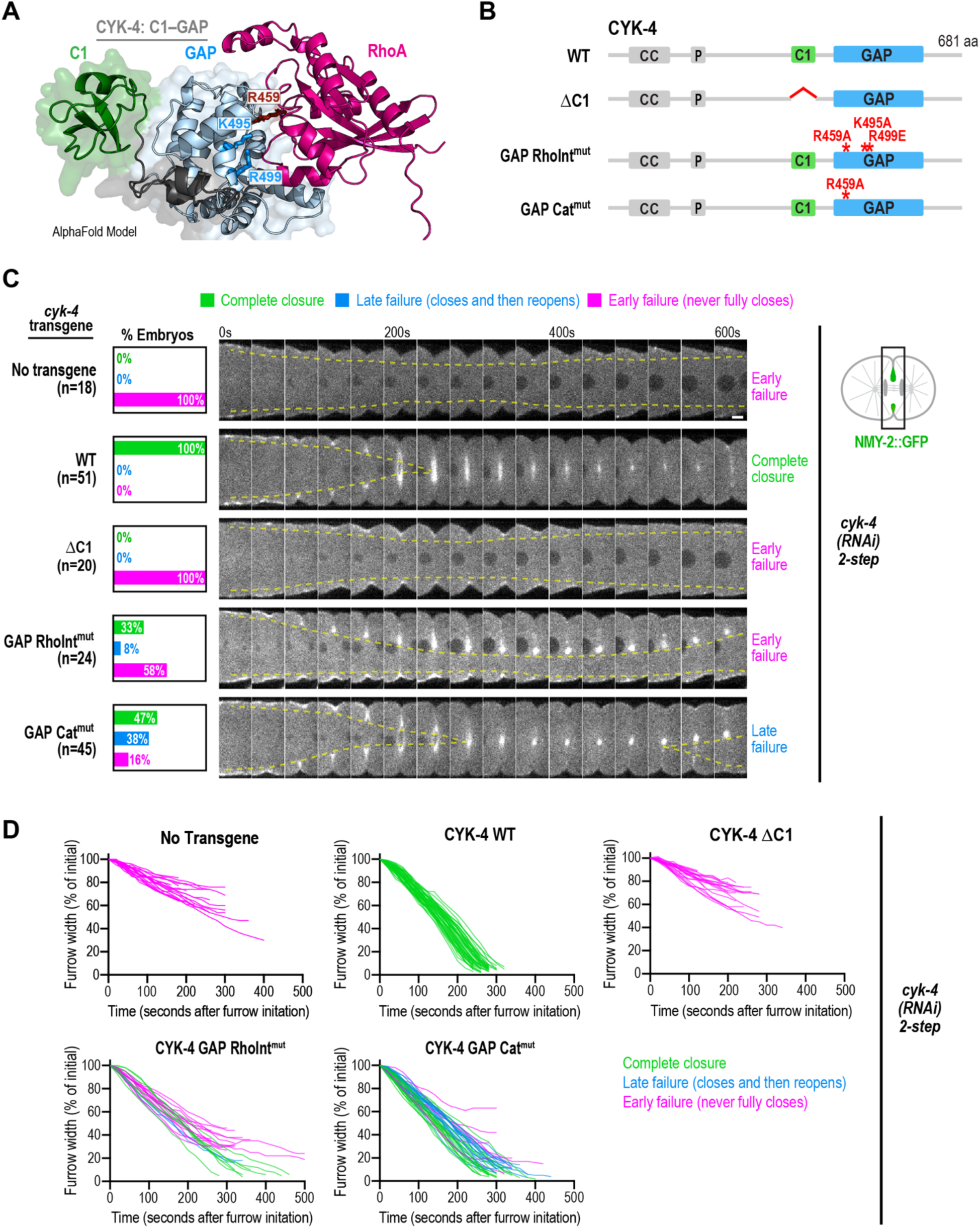
Distinct effects of disrupting CYK-4’s Rho-binding interface versus selective mutation of its GAP catalytic activity on cytokinesis. (**A**) AlphaFold2^16^ model showing the predicted interaction between the C1-GAP domain region of *C. elegans* CYK-4 and RhoA (encoded by *rho-1*). (**B**) Schematics illustrate the proteins encoded by a set of single-copy untagged RNAi-resistant transgenes under control of endogenous *cyk-4* regulatory sequences inserted into a specific site on chromosome 2. Two CYK-4 GAP domain mutants were analyzed: GAP RhoInt^mut^ disrupts the interface that allows the GAP domain to interact with Rho family GTPases and GAP Cat^mut^ is expected to prevent the GAP domain from catalyzing GTP hydrolysis without disrupting RhoA binding. Transgenes encoding WT CYK-4 and CYK-4 lacking its C1 domain (ΔC1) required for membrane association^5,10^ served as controls. (**C**) Strains expressing a GFP fusion with non-muscle myosin II (NMY-2::GFP) along with the indicated untagged *cyk-4* transgenes in the background of CYK-4::GFP::MEI-1N were treated using the two-step RNAi protocol described in *Figure S1F*. Images show the furrow region in representative timelapse sequences of embryos from the indicated conditions. Graphs to the left indicate the percentage of embryos that exhibited early failure (defined as failure of the furrow to fully close), late failure (defined as complete closure followed by reopening) or successfully completed cytokinesis for each condition. Semi-transparent dashed yellow lines track the inside edges of the furrow to facilitate visualization of closure phenotypes. (**D**) Plots of the kinetics of contractile ring closure in individual embryos for the conditions shown in *C*. Traces are color-coded to mark embryos that exhibited early failure (*magenta*), late failure (*blue*), or completed cytokinesis (*green*). Times are seconds after furrow initiation. Scale bar, 5 μm. *See also Figures S1-S3*.

Next, we focused on analyzing the engineered CYK-4 GAP domain mutants. GAP RhoInt^mut^ mutant CYK-4, while better at supporting cytokinesis than ΔC1 CYK-4, slowed the rate of furrow ingression compared to WT (**Fig. 1C,D**; **Fig. S3B,C**). Notably, cytokinesis failed in about two-thirds of GAP RhoInt^mut^ embryos, with many exhibiting an early failure in which the furrow never fully closed (58% of NMY-2::GFP and 27% of mCherry::plasma membrane & histone embryos) and the remainder exhibiting a late failure in which the furrow closed fully and then reopened (8% of NMY-2::GFP and 33% of mCherry::plasma membrane & histone embryos; **Fig. 1C,D**; **Fig. S3B,C**). The CYK-4 GAP activity mutant (GAP Cat^mut^) was significantly better at supporting cytokinesis than GAP RhoInt^mut^ (**Fig. 1C,D; Fig. S3B,C**). However, while the rate of furrow ingression in GAP Cat^mut^ embryos was similar to WT, they also exhibited significant rates of cytokinesis failure (54% and 21%, respectively, in the NMY-2::GFP and mCherry plasma membrane & histone backgrounds; **Fig. 1C,D**; **Fig. S3B,C**). In contrast to the prevalent early failure in GAP RhoInt^mut^ embryos, GAP Cat^mut^ embryos were strongly biased to late failure in which the furrow fully closed and then reopened later (**Fig. 1C,D**). Collectively, these results suggest that the ability of the CYK-4 GAP domain to bind Rho GTPases and the ability to promote GTP hydrolysis are both important for cytokinesis. However, the phenotypic differences between the two GAP domain mutants hint that some functions of the GAP domain may not require the ability to promote GTP hydrolysis.

### An assay for monitoring recruitment of centralspindlin to the equatorial cortex during cytokinesis

Next, we wanted to compare the ability of WT, GAP RhoInt^mut^, and GAP Cat^mut^ CYK-4 to facilitate the recruitment of centralspindlin to the equatorial cortex during cytokinesis. We employed the ΔC1 mutant, which is expected to prevent centralspindlin targeting, as a control. Measuring the cortical targeting of centralspindlin is challenging because of the rapid cell shape changes that occur during cytokinesis and because the amount of centralspindlin on the cortex is very low compared to the amount at the adjacent central spindle (**Fig. 2A**, white compared to green arrowheads in control). To overcome these limitations, we *in situ* tagged ZEN-4, the kinesin-6 subunit of centralspindlin, with mScarlet and analyzed its recruitment in embryos depleted of NMY-2 to prevent cell shape changes and SPD-1, the *C. elegans* homolog of the microtubule bunding PRC1, to disrupt central spindle assembly (**Fig. 2A,B**). To detect ZEN-4 on the equatorial cortex, we collected a cortical z-stack and generated a maximum intensity projection of the three z-slices closest to the coverslip over the three time points between 260 and 300s after anaphase onset (**Fig. 2B**). As contractile rings during the first embryonic division have an asymmetric structure and close asymmetrically within the division plane^18–20^, equatorial ZEN-4 concentrates asymmetrically on one side of the embryo. Thus, depending on the rotational orientation of the embryo relative to the coverslip, the equatorial ZEN-4 signal in the maximum intensity projection of the coverslip-adjacent cortex could span all or part of the equator or be absent when ZEN-4 was concentrated on the side of the embryo away from the coverslip. We quantified the ZEN-4 signal by performing a wide line scan drawn parallel to the embryo’s long axis and centered over visible cortical ZEN-4 signal, or over the central axis of the embryo in cases with no detectable cortical ZEN-4 signal (**Fig. 2B**). We started by analyzing mScarlet::ZEN-4 recruitment in embryos expressing transgenic untagged WT or ΔC1 mutant CYK-4 in the presence of endogenous CYK-4. In embryos expressing transgenic WT CYK-4, we observed a strong accumulation of mScarlet::ZEN-4 on the equatorial cortex. By contrast, the equatorial recruitment of mScarlet::ZEN-4 was almost completely suppressed by the presence of the transgenic CYK-4 ΔC1, even when endogenous CYK-4 was not depleted (**Fig. 2C,D**). Immunoblotting of worms expressing untagged transgenic WT and ΔC1 CYK-4 has shown that they are expressed at levels equivalent to those of endogenous CYK-4^12^. Since centralspindlin is a heterotetramer that contains two molecules of CYK-4 and two molecules of ZEN-4, we suspect that ∼25% of the centralspindlin contains two molecules of WT CYK-4, ∼25% contains two molecules of ΔC1 mutant CYK-4 and ∼50% contains one molecule of WT CYK-4 and one molecule of ΔC1 mutant CYK-4. The strong suppression of mScarlet::ZEN-4 recruitment suggests that centralspindlin complexes require both of their C1 domains to engage with the plasma membrane and be recruited to the cortex and that the recruitment of complexes containing even one molecule of ΔC1 CYK-4 is compromised. Overall, these results highlight that the approach of preventing furrow ingression (via NMY-2 depletion) and centralspindlin concentration on the spindle midzone (via SPD-1 depletion), in conjunction with imaging of *in situ*-tagged ZEN-4, enables robust monitoring of centralspindlin recruitment to the equatorial cortex.

**Figure 2.**
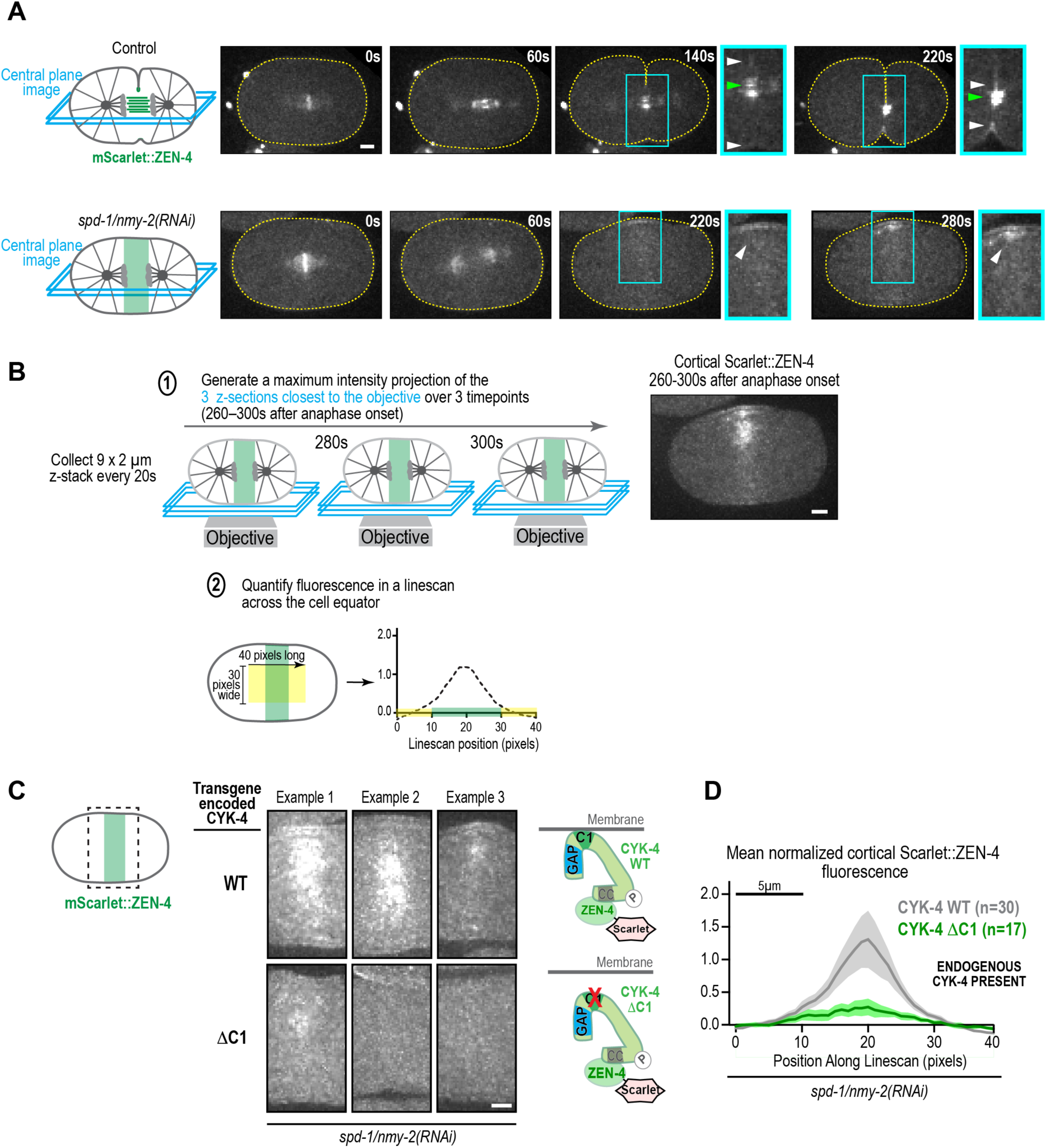
An assay for monitoring the recruitment of centralspindlin to the equatorial cortex during cytokinesis. (**A**) Representative central plane images of embryos expressing an mScarlet fusion with the centralspindlin component ZEN-4. (*top row*) Monitoring centralspindlin recruitment to the equatorial cortex in control embryos is challenging because of cell shape changes during cytokinesis (*indicated by dashed yellow outlines*) and by the proximity of the dim cortical centralspindlin signal (*white arrowheads*) to the bright signal on the microtubule bundles of the central spindle (*green arrowheads*). (*bottom row*) In our assay condition (*spd-1/nmy-2(RNAi)*), non-muscle myosin II was depleted to prevent cell shape changes and the *C. elegans* PRC1 homolog SPD-1 was depleted to disrupt the microtubule bundles of the central spindle. In these embryos, centralspindlin accumulates on the cortex in an equatorial band. Times are seconds after anaphase onset. Cyan boxed regions are magnified 1.5X in the adjacent images. (**B**) Schematics illustrate how the centralspindlin on the equatorial cortex was quantified. Maximum intensity projections were generated of the 3 z-planes closest to the coverslip surface, which contained the cortex, over the three timepoints between 260 and 300s after anaphase onset, and a linescan was performed to quantify the equatorial signal (see methods). (**C**) Three representative examples of the cortical accumulation of mScarlet::ZEN-4 in embryos expressing WT (*top*) or ΔC1 CYK-4 (*bottom*) in the presence of endogenous CYK-4. Schematics to the right illustrate the two constructs. (**D**) Graphs plot mean normalized cortical mScarlet::ZEN-4 fluorescence as a function of position along the linescan for each condition. The shaded region for each curve represents the 95% CI. Scale bars on the images and graph, 5μm.

### Interaction of the CYK-4 GAP domain with RhoA promotes the recruitment of centralspindlin to the equatorial cortex during cytokinesis

Since the GAP domain is adjacent to the C1 domain in the C-terminal half of CYK-4 (**Fig. 1A,B**), we next tested if the GAP domain cooperates with the C1 domain to facilitate the cortical recruitment of centralspindlin by interacting with RhoA-GTP in the plasma membrane (**Fig. 3A**). We compared mScarlet::ZEN-4 recruitment in embryos expressing transgenic untagged WT CYK-4 and GAP RhoInt^mut^ CYK-4 in the presence of endogenous CYK-4. As was the case for CYK-4 ΔC1, the equatorial recruitment of mScarlet::ZEN-4 was strongly suppressed by the presence of the transgene-encoded GAP RhoInt^mut^ CYK-4, even when endogenous CYK-4 was not depleted (**Fig. 3B,C**). The cortical recruitment of mScarlet::ZEN-4 was reduced even further when endogenous CYK-4 was depleted to make GAP RhoInt^mut^ the only source of CYK-4 (**Fig. 3D,E**). By contrast, the equatorial recruitment of mScarlet::ZEN-4 was not reduced when GAP Cat^mut^ CYK-4 was expressed in the presence or absence of endogenous CYK-4 (**Fig. 3B-E**), indicating that GAP domain catalytic activity is not essential for promoting the cortical recruitment of centralspindlin. To determine if the CYK-4 GAP domain promotes centralspindlin recruitment by binding to RhoA, we compared ZEN-4 recruitment in SPD-1 depleted embryos in which furrow ingression was prevented by depletion of RhoA (*rho-1(RNAi)*) to embryos in which furrow ingression was prevented by depletion of non-muscle myosin II (*nmy-2(RNAi)*). Depletion of RhoA suppressed recruitment of mScarlet::ZEN-4 as effectively as the GAP RhoInt^mut^ CYK-4 mutant (**Fig. 3F**). Collectively, these results suggest that the Rho-binding interface of the CYK-4 GAP domain is required for the recruitment of centralspindlin to the equatorial cortex during cytokinesis, whereas the GAP domain’s catalytic activity is not.

**Figure 3.**
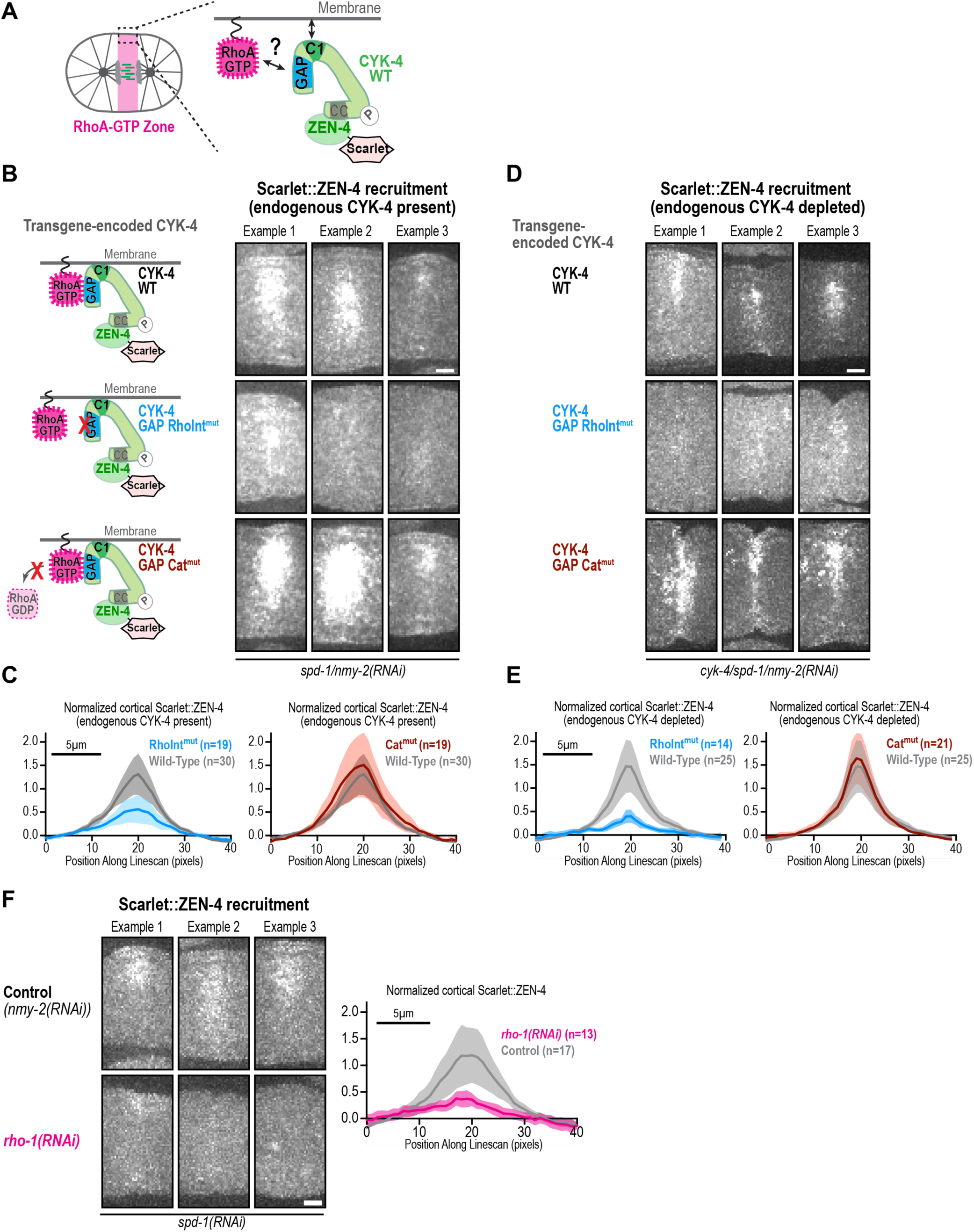
The CYK-4 GAP Rho-binding interface promotes the cortical recruitment of centralspindlin. (**A**) Schematic illustrating the possibility tested here that interaction of the CYK-4 GAP domain with active GTP-bound RhoA could collaborate with the C1 domain to facilitate recruitment of centralspindlin to the equatorial cortex during cytokinesis. (**B, D**) Three representative examples of the cortical accumulation of mScarlet::ZEN-4 in embryos expressing WT (*top*), GAP RhoInt^mut^ (*middle*) or GAP Cat^mut^ (*bottom*) CYK-4 with endogenous CYK-4 present (*B*) or absent (*D*). Schematics on the left in (*B*) illustrate the three constructs. CYK-4 WT examples in *B* are reproduced from Figure 2C for comparison. (**C, E**) Graphs plot mean normalized cortical mScarlet::ZEN-4 fluorescence as a function of position along the linescan for each condition. (**F**) Three representative examples of the cortical accumulation of mScarlet::ZEN-4 in control (*top*) and RhoA depleted (*rho-1(RNAi), bottom)* embryos. Graph on the right plots mean normalized cortical mScarlet::ZEN-4 fluorescence as a function of position along the linescan for each condition. The shaded region for the curves in the graphs represents the 95% CI. Scale bars on the images and graph, 5μm.

### The CYK-4 GAP Rho-Interaction and catalytic mutants have opposite effects on furrowing when Rho activation is partially compromised

In our analysis of furrow ingression, GAP RhoInt^mut^ and GAP Cat^mut^ embryos both experienced elevated rates of cytokinesis failure, although GAP RhoInt^mut^ embryos tended to exhibit slowed furrowing and early failure, whereas GAP Cat^mut^ embryos tended to exhibit late failure or succeed (**Fig. 1C,D**; **Fig. S3B,C**). Our analysis of centralspindlin recruitment suggests that these phenotypes may have distinct origins. GAP RhoInt^mut^ embryos might fail cytokinesis due to a reduced ability to recruit centralspindlin to the cortex, which would limit RhoA activation. By contrast, GAP Cat^mut^ embryos do not have a defect in centralspindlin recruitment and might fail cytokinesis for a different reason. If this were true, then GAP RhoInt^mut^ embryos should be more sensitive to a perturbation that reduces the levels of active RhoA than GAP Cat^mut^ embryos. To test this prediction, we took advantage of the *nop-1*(*it142*) mutant, which disrupts a non-essential nematode-specific protein that functions to promote RhoA activation in parallel to centralspindlin^17^. In the centralspindlin recruitment assay, mScarlet::ZEN-4 levels are modestly reduced in *nop-1(it142)* embryos, consistent with the expected reduction in active RhoA-GTP (**Fig. 4A-C**). Notably, in agreement with the prediction above, furrowing was severely compromised when *nop-1(it142)* was combined with GAP RhoInt^mut^ CYK-4. In contrast, furrowing was not impaired when *nop-1(it142)* was combined with GAP Cat^mut^ CYK-4 (**Fig. 4D**). These results provide functional support for the conclusion that the Rho-binding interface of the CYK-4 GAP domain promotes recruitment of centralspindlin to the equatorial cortex, whereas the catalytic activity of the GAP domain is not essential for centralspindlin recruitment.

**Figure 4.**
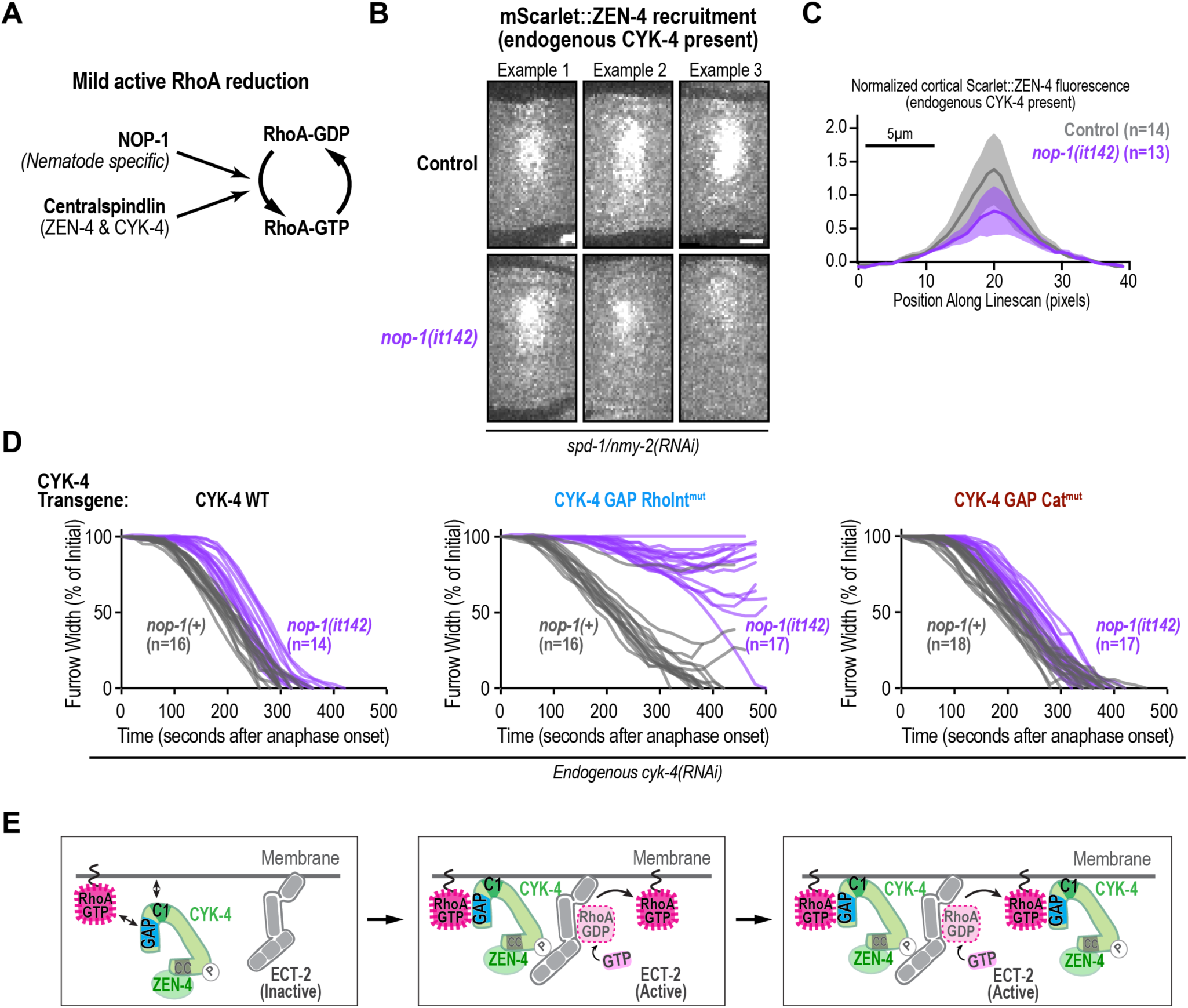
The CYK-4 GAP Rho-Interaction and catalytic mutants have opposite effects on furrowing when Rho activation is partially compromised. (**A**) Schematic illustrating the previously proposed model that NOP-1 functions in parallel to centralspindlin to promote RhoA activation^17^. Although NOP-1 contributes to contractility and slightly accelerates cytokinesis onset, it is not essential for cytokinesis. (**B**) Three representative examples of the cortical accumulation of mScarlet::ZEN-4 in control (*top*) and *nop-1(it142)* (*bottom*) embryos. (**C**) Graph plots mean normalized cortical mScarlet::ZEN-4 fluorescence as a function of position along the linescan for each condition. The shaded region for the curves in the graphs represents the 95% CI. (**D**) Plots of the kinetics of contractile ring closure in control *nop-1(+)* (*grey*) and *nop-1(it142)* (*purple*) embryos expressing WT, GAP RhoInt*^mut^*, and GAP Cat^mut^ CYK-4 after depletion of endogenous CYK-4. (**E**) Model showing how recruitment of centralspindlin to the cortex via an interaction between the CYK-4 GAP domain and RhoA–GTP at the membrane generates positive feedback in the circuit that generates the equatorial RhoA zone during cytokinesis. Interactions between the CYK-4 C1 and GAP domains with membrane and membrane-anchored active RhoA (RhoA-GTP) recruit the centralspindlin complex to the cell surface, where CYK-4 can then activate the RhoGEF ECT-2, leading to the generation of more active RhoA that can recruit more centralspindlin. Scale bars, 5μm.

We note that our results are in contrast to a prior report that suggested that replacing CYK-4 with a GAP Cat mutant leads to a complete suppression of furrowing in the *nop-1(it142)* background^10^. We confirmed the presence of the *nop-1(it142)* and GAP Cat mutations by sequencing to exclude a trivial explanation for this difference. One possible reason for the difference is that we are employing untagged *cyk-4* transgenes expressed under their own regulatory sequences. In contrast, the prior study expressed C-terminally GFP-tagged CYK-4 using the *cyk-4* promoter and *pie-1* 3’UTR, which might lead to differences in protein functionality and expression that synergize with the effects of GAP Cat^mut^ CYK-4 (see section below).

Collectively, these results highlight a major difference between the effects of disrupting the Rho-binding interface of the CYK-4 GAP domain and disrupting its catalytic activity. The severe defect observed when the Rho-binding interface was disrupted in *nop-1(it142)* embryos provides strong support for a positive feedback model in which RhoA-GTP promotes the recruitment of centralspindlin, which, in turn, activates ECT-2 to drive the generation of more RhoA-GTP (**Fig. 4E**).

### The catalytic activity of the CYK-4 GAP domain releases centralspindlin from the cortex

Our results above suggested that the primarily late cytokinesis failure we observe in GAP Cat^mut^ embryos in the presence of wild-type *nop-1* (**Fig. 1C,D**; **Fig. S3B,C**) is not due to a failure to recruit centralspindlin, suggesting that GAP catalytic activity contributes a distinct function. In the cortical recruitment assay, where no furrow ingression occurs, and the central spindle is absent, we observed that mScarlet::ZEN-4 was eventually largely released from the cortex, whereas GAP Cat^mut^ CYK-4, mScarlet::ZEN-4 persisted on the cortex until the end of imaging (*not shown*). This chance observation suggested that GAP domain catalytic activity may be important to release centralspindlin from the cortex (**Fig. 5A**). To directly assess the effect of disrupting GAP catalytic activity on the release of centralspindlin from the cortex, we collected a new dataset using the recruitment assay conditions (mScarlet::ZEN-4; NMY-2, SPD-1, and endogenous CYK-4 depleted) and imaging the embryos for ∼800s after anaphase onset so that we could monitor loss of cortical centralspindlin. We imaged 38 WT CYK-4 embryos and 37 CYK-4 GAP Cat^mut^ embryos and analyzed loss of cortical centralspindlin in embryos that exhibited no furrowing (**Fig. S4A-D**) or <50% furrowing (**Fig. 5B,C**, **Fig. S4D**), which yielded similar results (we note that compared to embryos expressing WT CYK-4, it was more difficult to obtain GAP Cat^mut^ CYK-4 embryos that exhibited no furrowing; **Fig. S4A**). In embryos expressing WT CYK-4, the cortical centralspindlin signal (mScarlet::ZEN-4) tended to increase and then dissipate, with the majority of the signal gone by ∼500s after anaphase onset (**Fig. 5B**, **Fig. S4D**). In embryos expressing GAP Cat^mut^ CYK-4, cortical mScarlet::ZEN-4 remained at the cortex, typically oscillating in intensity, over the full 800s window, often reaching levels higher than those measured in the 260-300s time window used for our single timepoint recruitment assay (**Fig. 5C**, **Fig. S4D**). We conclude that while the GAP activity of CYK-4 is not required to recruit centralspindlin to the cortex, it is required to facilitate its release from the cortex. It is worth noting that since our recruitment data suggest that both GAP domains need to be able to bind RhoA GTP to efficiently target centralspindlin to the cortex, hydrolysis by one of the two GAP domains in a tetramer might be sufficient to release centralspindlin from the cortex.

**Figure 5.**
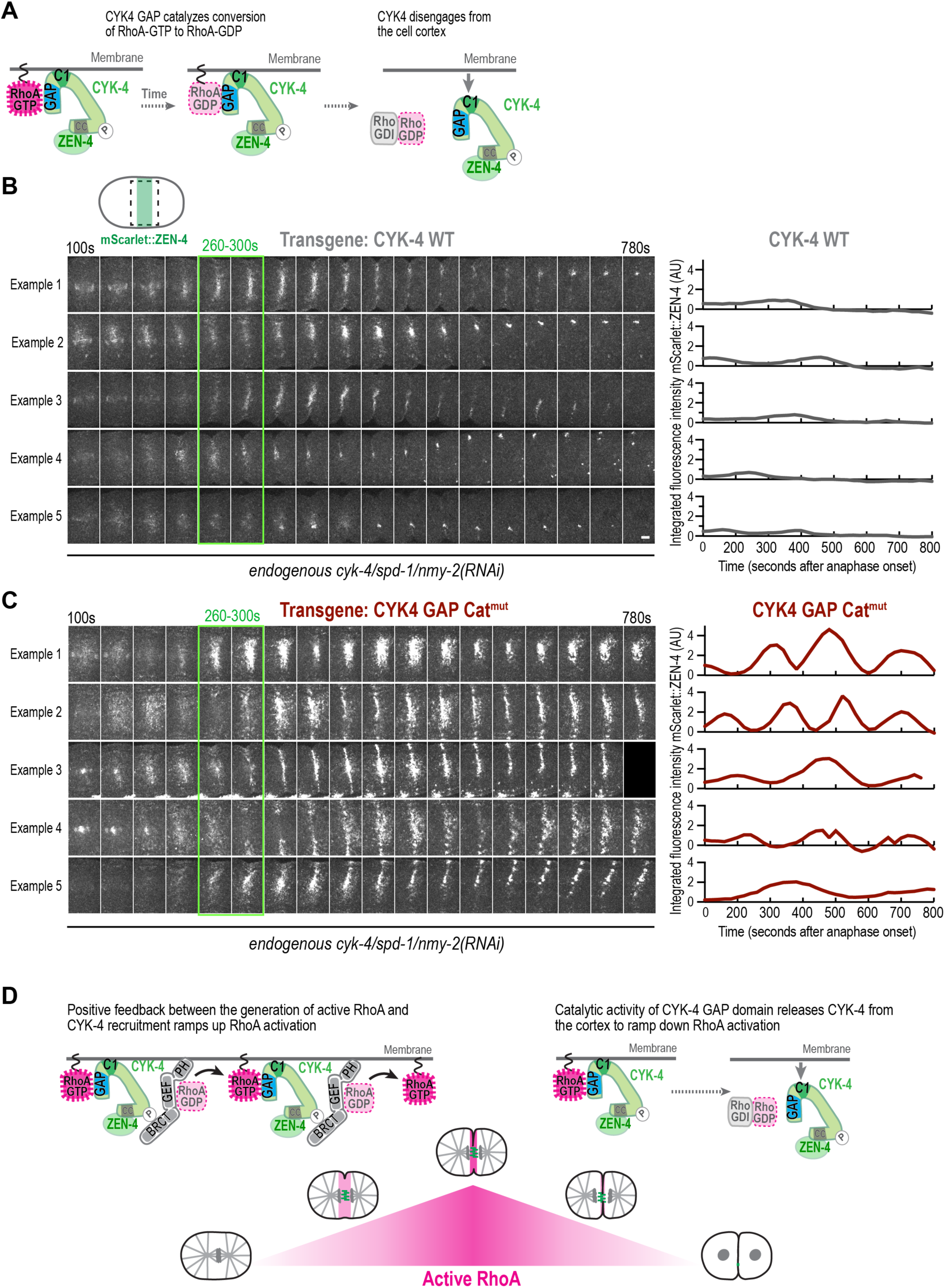
The CYK-4 GAP catalytic activity serves as an intrinsic timer to limit the cortical localization of centralspindlin. (**A**) Schematic showing how the catalytic activity of the CYK-4 GAP domain could release CYK-4 from the surface by converting RhoA-GTP to RhoA-GDP. (**B, C**) (*left*) Images of the furrow region from representative timelapse sequences of embryos expressing WT (**B**) or GAP Cat^mut^ CYK-4 (**C**). Images are maximum intensity projections of full embryo z-stacks. (*right*) Quantification of total fluorescence intensity for the images on the left. The example images and quantification shown are of the five embryos with the highest signal that exhibited <50% furrowing for both conditions. A montage of all other embryos exhibiting no or <50% furrowing with discernable signal is shown in *Figure S4A-C* along with a graphical summary of extent of furrowing and presence of signal for all embryos imaged. Times are seconds after anaphase onset. Green boxes mark the time interval used for the centralspindlin recruitment assay described in *Figure 2B,C*. (**D**) Model showing how the CYK-4 GAP domain could contribute to the positive feedback that generates the equatorial zone of active RhoA that sustains the contractile ring (*left*) and how the catalytic activity of the CYK-4 GAP domain allows centralspindlin to be released from the cell surface, which allows cortical activity to ramp down as cytokinesis completes (*right*). *See also Figure S4*.

Extensive recent work has highlighted the ability of Rho GTPases to form dynamic self-organized zones that arise from complex networks of positive and negative feedback^1^. Here, we elucidate a key element of the positive feedback that leads to the rapid formation of an equatorial RhoA zone that drives robust cytokinesis. We show that the CYK-4 GAP domain employs its RhoA-binding interface to recruit centralspindlin to the cortex. Cortical centralspindlin, in turn, activates ECT-2 to drive the generation of more RhoA-GTP (**Fig. 5D**). Thus, centralspindlin binds to active RhoA and leads to the generation of more active RhoA. Our results further suggest that the catalytic activity of the CYK-4 GAP domain serves as an intrinsic timer that may limit the residence time of centralspindlin at the cortex.

Our work suggests that the recruitment of centralspindlin to the cell cortex depends on the engagement of both of its GAP domains with RhoA-GTP. While centralspindlin is at the cortex, it also interacts with and activates ECT2, which requires an interaction between the CYK4 N-terminus and the ECT2 BRCT domain^22–26^. In addition to the recruitment of centralspindlin via an interaction of the CYK-4 GAP domain with RhoA-GTP that we describe here, a second source of positive feedback that has been proposed to contribute to RhoA generation during cytokinesis is binding of RhoA-GTP to an allosteric site on the ECT2 PH domain (independent of RhoA binding to the catalytic site of its GEF domain). Binding of RhoA-GTP to the ECT2 allosteric site is thought to facilitate ECT2 activation by preventing ECT2’s PH domain from occluding its GEF active site^21^. While these two sources of positive feedback could act separately during RhoA activation and cytokinesis, it is attractive to speculate that the RhoA-GTP that interacts with the CYK4 GAP domain to recruit centralspindlin to the cortex is the same RhoA-GTP that binds to the allosteric site of ECT2 to activate it. For example, centralspindlin may be recruited to the cortex via interaction of its GAP domain with RhoA-GTP and then the RhoA-GTP–CYK4 complex could engage in a two-point interaction with ECT2 to activate it in which the PLK1-phosphorylated CYK4 N-terminus interacts with the ECT2 BRCT domain^22–26^ while the RhoA-GTP bound to the CYK4 GAP domain interacts with the ECT2 allosteric site. Future biochemical and structural work will be important to explore this and other possibilities.

## ACKNOWLEDGEMENTS

The authors thank Rebecca Green for help with the model figure and Esther Zanin for the suggestion to more carefully consider the effects of GAP Cat^mut^ CYK-4 in the centralspindlin recruitment assay. This work was supported by grants from the NIH to K.O. (R01 GM147265) and A.J.S. (F32 GM145068). K.O. and A.D. acknowledge partial salary support from the Ludwig Institute for Cancer Research.

## AUTHOR CONTRIBUTIONS

Conceptualization: K.O., A.J.S, K-Y.L., J.S.G-C.; Methodology: A.J.S, K-Y.L., J.S.G-C., K.O.; Formal analysis: K-Y.L., A.J.S.; Investigation: A.J.S, K-Y.L., J.S.G-C., P. L-G.; Resources: K.O., A.D.; Writing - original draft: A.J.S., A.D., K.O.; Writing - review & editing: A.J.S., A.D., K.O.; Visualization: A.J.S, K.O.; Supervision: A.D., K.O.; Project administration: A.D., K.O.; Funding acquisition: A.J.S, A.D., K.O

## SUPPLEMENTAL FIGURE TITLES AND LEGENDS

**Figure S1.**
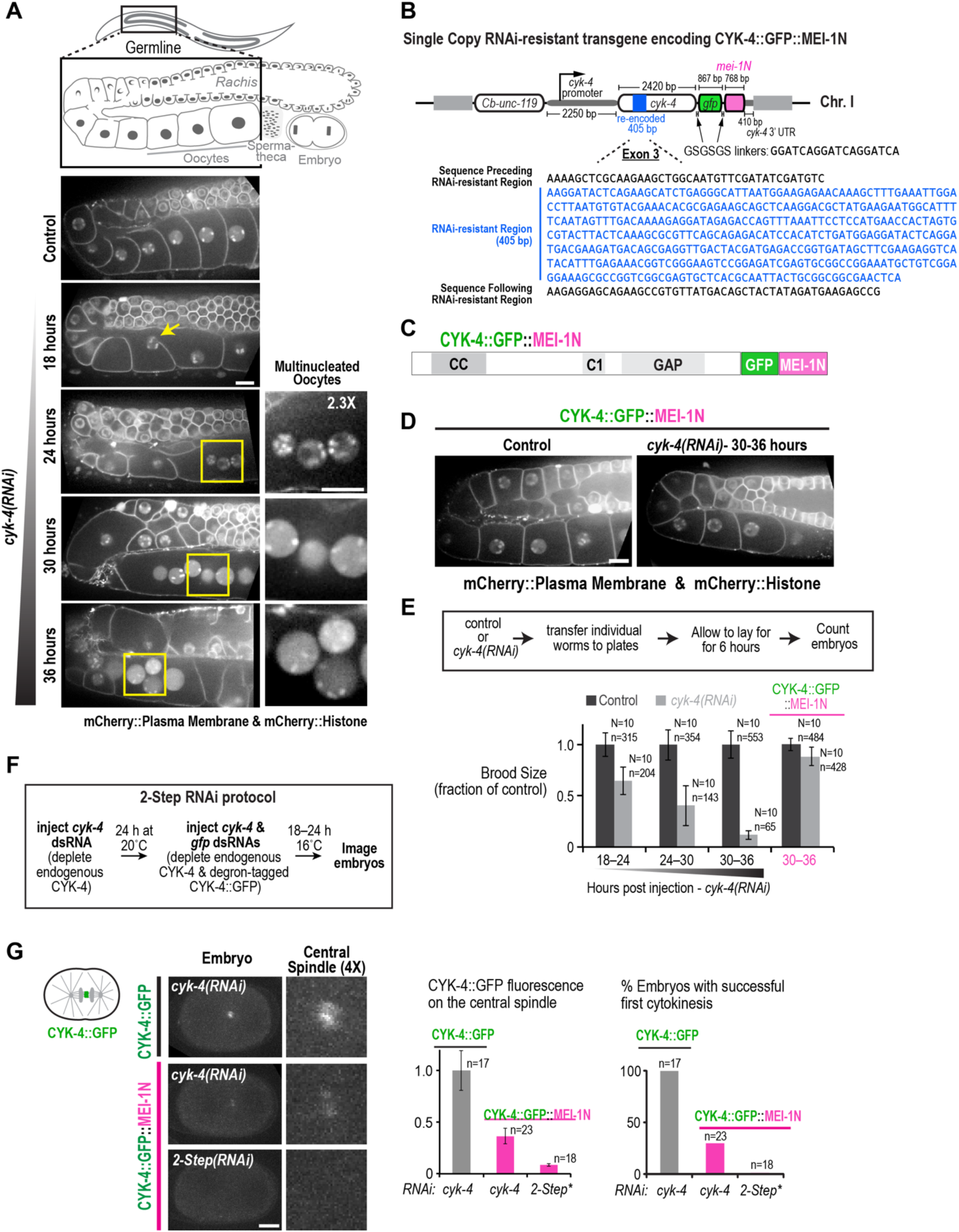
A *2-step RNAi* approach to bypass the essential role of CYK-4 in the germline all allow robust depletion of CYK-4 in one-cell stage *C. elegans* embryos (related to Figure 1). (**A**) Hermaphrodites expressing mCherry-tagged plasma membrane and histone probes were injected with dsRNA targeting *cyk-4*. Worms were anesthetized 18, 24, 30, and 36 hours post-injection as indicated (n=5 for each timepoint), and the germline region (see schematic) was imaged. Representative images illustrate the progressive defects in germline structure that become more severe as the CYK-4 protein is depleted. Yellow arrow marks a nucleus falling out of a forming oocyte. Yellow boxes highlight multinucleated regions adjacent to the spermatheca, where compartmentalized oocytes should normally be, shown at higher magnification in the adjacent panels. (**B**) Schematic detailing the RNAi-resistant single-copy transgene that encodes CYK-4 tagged with GFP and the N-terminal 224 amino acids of MEI-1 (*degron, pink*), inserted into a locus on Chromosome I^27^. The genomic sequence from the *cyk-4* locus including 2250bp upstream of the start codon and 410 bp downsteam of the stop codon was modified by re-encoding 405 bp in exon 3 without altering the protein sequence to render the transgene resistant to a dsRNA targeting endogenous *cyk-4*; a sequence encoding GFP and the first 224 aa of MEI-1 was inserted before the 3’UTR. (**C**) Schematic of the protein encoded by the transgene in *B*. (**D**) Representative images of the germline region in hermaphrodites expressing mCherry-tagged plasma membrane and histone probes and CYK-4::GFP::MEI-1N. The germline in control (uninjected) worms and worms injected with *cyk-4* dsRNA were imaged after 30-36 hours (n=5). (**E**) Worms expressing the mCherry-tagged plasma membrane and histone probes, without or with the transgene encoding CYK-4::GFP::MEI-1N as indicated, were injected with *cyk-4* dsRNA (*grey bars*) or left uninjected as a control (*black bars*). Brood size was scored by counting the total number of progeny laid within the indicated 6-hour post-injection time windows (N is the number of worms, n is total embryos counted, error bars are the SD). Brood size decreased over time in worms injected with *cyk-4* dsRNA that lacked the CYK-4::GFP::MEI-1N transgene, but was maintained in worms with the transgene. (**F**) Outline of the two-step RNAi protocol that enables robust CYK-4 depletion. (**G**) Worms expressing CYK-4::GFP::MEI-1N, or CYK-4::GFP (lacking the MEI-1N tag) as a control, were treated using a single-step *cyk-4* RNAi protocol. Images on the left, quantified in the graph in the middle, show that the CYK-4::GFP::MEI-1N signal at the central spindle is much lower than the signal for CYK-4::GFP lacking the MEI-1N degron. The graph on the right shows that cytokinesis failed in ∼75% of cases in embryos expressing the degron-tagged protein but not in embryos expressing CYK-4::GFP. In embryos with the CYK-4::GFP::MEI-1N transgene subjected to the 2-step protocol, the CYK-4::GFP::MEI-1N signal at the central spindle was reduced below the level of detection, and 100% of embryos failed cytokinesis. Error bars are SEM. Scale bars,10µm.

**Figure S2.**
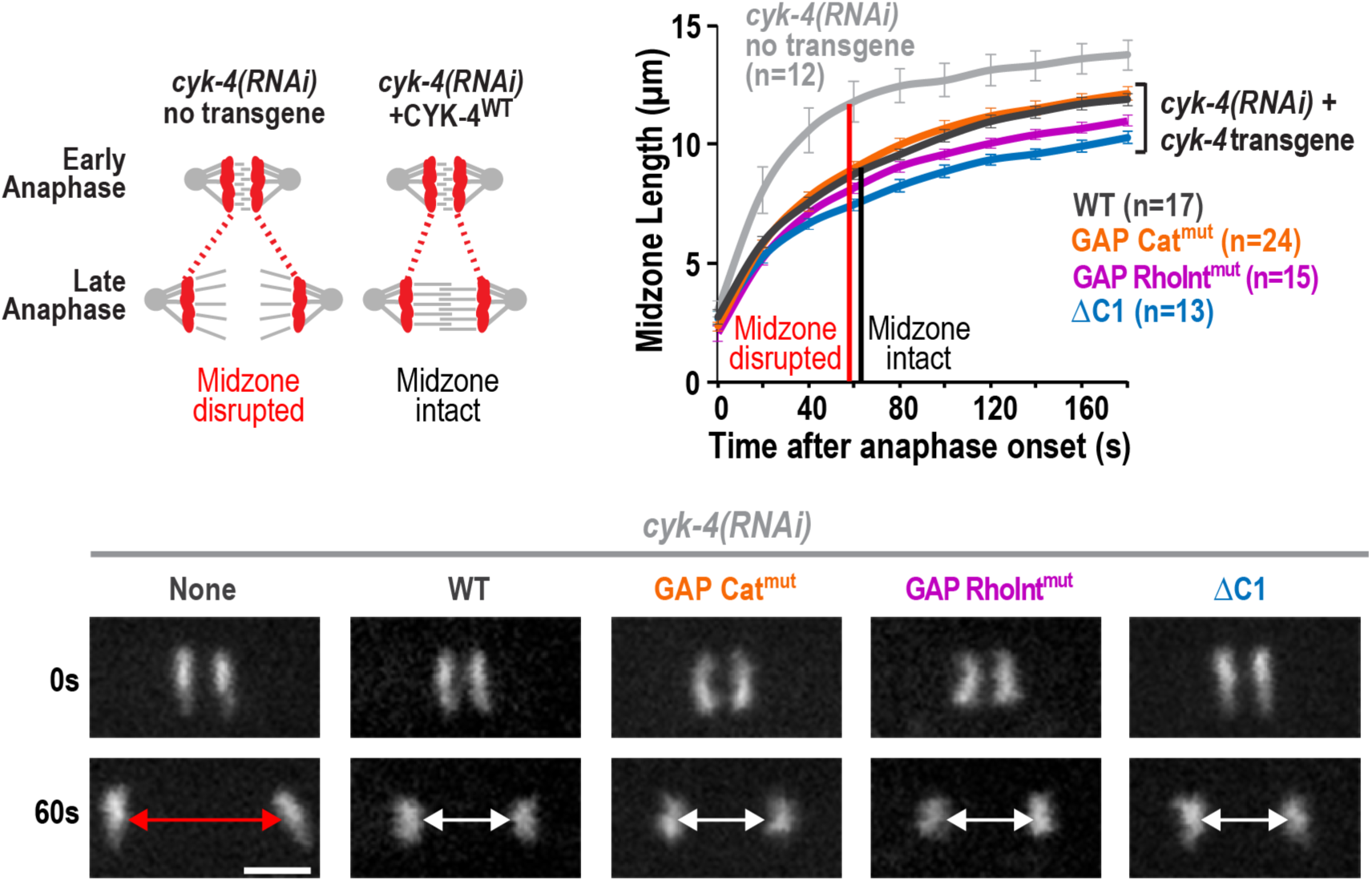
Engineered CYK-4 variants do not disrupt central spindle assembly (Related to Figure 1). Strains expressing untagged CYK-4 variants in the background of CYK-4::GFP::MEI-1N were treated using the two-step RNAi protocol described in Figure 1F. Graph plots the distance between the segregating sister chromosomes marked by mCherry::Histone. Times are seconds after anaphase onset. Error bars are the SEM. Scale bar, 5 μm.

**Figure S3.**
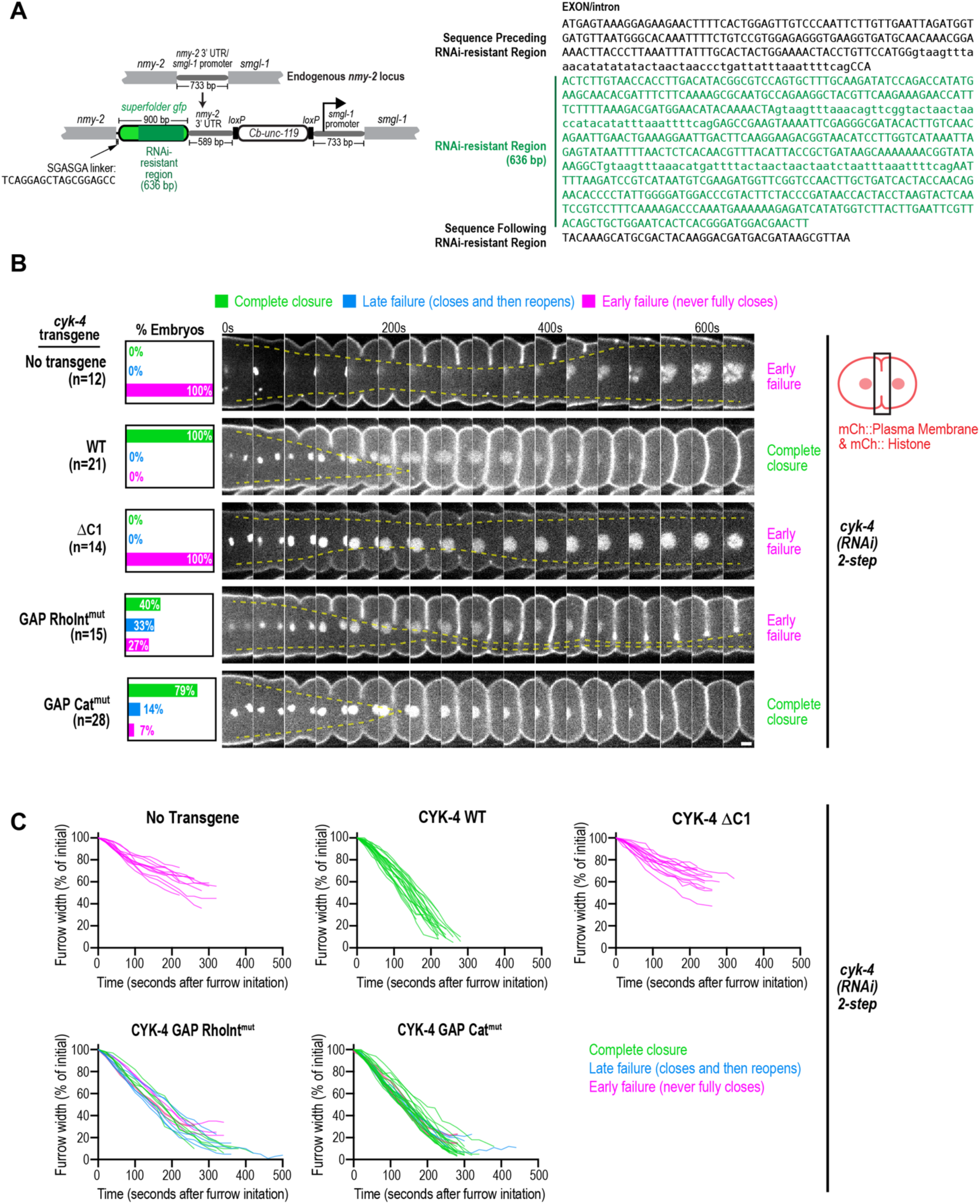
CRISPR-mediated tagging of NMY-2 with RNAi-resistant GFP and characterizing the effects of disrupting CYK-4’s Rho-binding interface versus GAP catalytic activity in embryos expressing mCherry-tagged plasma membrane and histone probes (Related to Figure 1). (**A**) Schematic (*left*) and sequence (*right*) detailing the RNAi-resistant GFP inserted to tag NMY-2 at its endogenous locus using CRISPR. (**B**) Strains expressing mCherry-tagged plasma membrane and histone probes along with the indicated untagged *cyk-4* transgenes in the background of CYK-4::GFP::MEI-1N were treated using the two-step RNAi protocol described in Figure S1F. Images show the furrow region in representative timelapse sequences of embryos from the indicated conditions. Graphs to the left indicate the percentage of embryos that exhibited early failure (defined as failure of the furrow to fully close), late failure (defined as complete closure followed by reopening) or successfully completed cytokinesis for each condition. Semi-transparent dashed yellow lines track the inside edges of the furrow to facilitate visualization of closure phenotypes. (**C**) Plots of the kinetics of contractile ring closure in individual embryos for the conditions shown in *C*. Traces are color-coded to mark embryos that exhibited early failure (*magenta*), late failure (*blue*), or completed cytokinesis (*green*). Times are seconds after furrow initiation. Scale bar, 5 μm.

**Figure S4.**
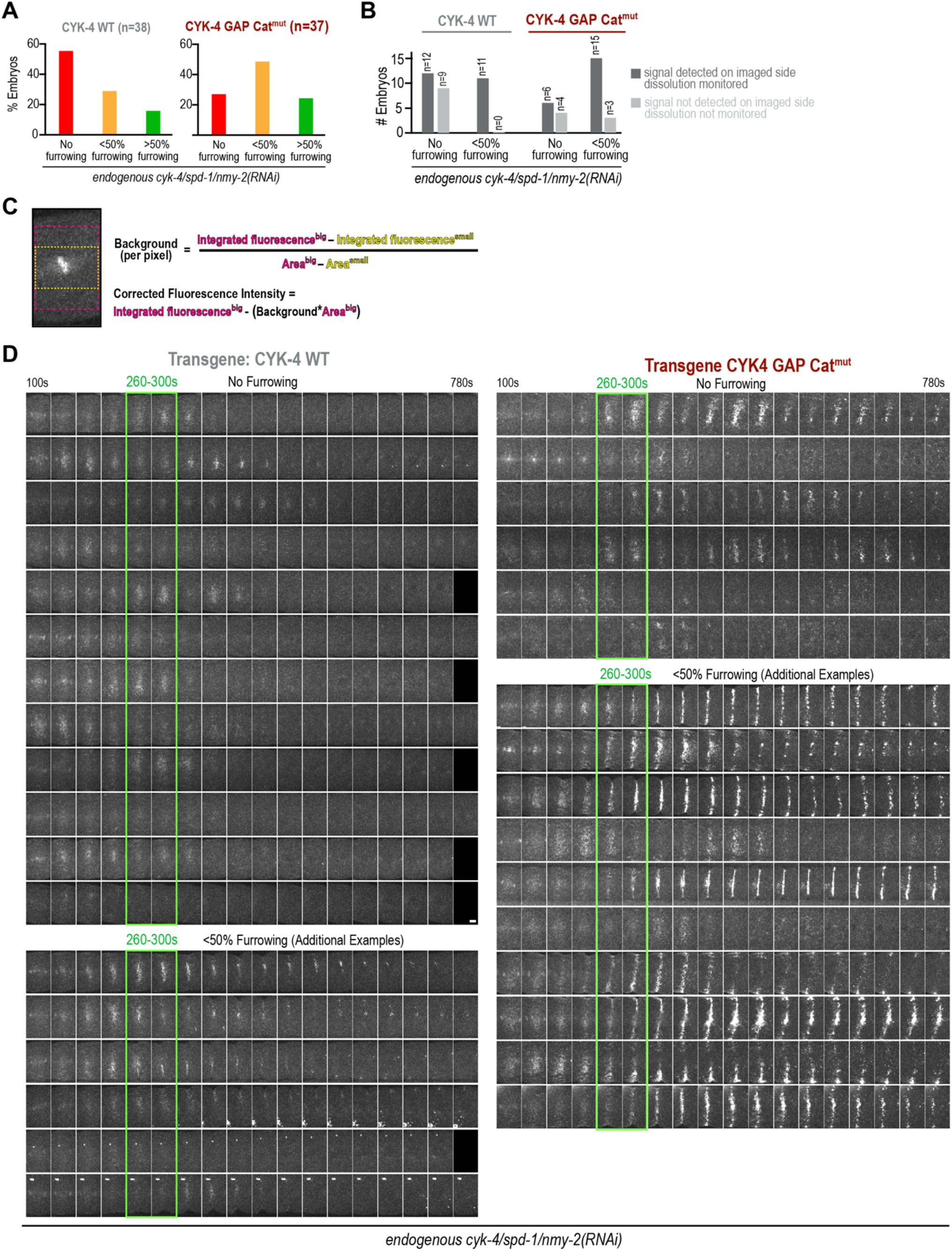
Additional information on analysis of dissolution of cortically localized centralspindlin (Related to Figure 5). (**A**) Graph plotting the percent of embryos undergoing “no furrowing”, “<50% furrowing”, or “>50% furrowing” among the embryos expressing transgenic WT or GAP Cat^mut^ CYK 4 after depletion of NMY-2, SPD-1, and endogenous CYK-4 that were imaged for the dissolution analysis shown in Figure 5B and below in *D*. (**B**) Graph showing the number of embryos that exhibited no furrowing or <50% furrowing that exhibited signal (*dark grey*) or did not exhibit signal (*light grey*) likely because of embryo orientation. Timelapse series for all embryos exhibiting signal are shown either in *D* or in Figure 5B. (**C**) Images and equations show how the integrated fluorescence intensity of mScarlet::ZEN-4 was quantified over time for the graphs in Figure 5B. (**D**) Additional examples of mScarlet::ZEN-4 recruitment in embryos expressing transgenic WT or GAP Cat^mut^ CYK 4 after depletion of NMY-2, SPD-1, and endogenous CYK-4 that either did not furrow (“no furrowing”) or experienced some furrowing (“<50% furrowing”). Images are maximum intensity projections of full embryo z-stacks. Times are seconds after anaphase onset. Green boxes mark the time interval used for the centralspindlin recruitment assay described in *Figure 2B,C*. Scale bars, 5μm.

## MATERIALS AND METHODS

### RESOURCE AVAILABILITY

#### Lead contact

Further information and requests for resources and reagents should be directed to and will be fulfilled by the lead contact, Karen Oegema.

#### Materials availability

All strains and other reagents generated in this study are freely available from the lead contact upon request.

#### Data and code availability

All primary data associated with the paper is available upon request.

### EXPERIMENTAL MODEL AND SUBJECT DETAILS

*C. elegans* strains (see Key Resources Table) were maintained at 20°C unless otherwise specified on standard nematode growth medium plates seeded with OP50-1 *E. coli*.

### METHOD DETAILS

#### C. elegans genome editing

Single-copy transgenes were generated using the transposon-based Mos single-copy insertion (MosSCI) method^27^. The single-copy transgene expressing CYK-4::GFP::MEI-1N was generated by cloning the sequence of interest (detailed in **Fig. S1B**) into pCFJ352 to create a transgene repair template. The pCFJ352-derived repair template was microinjected into EG6701, which has a Mos transposon insertion in the ttTi4348 site of chromosome I. The pCFJ352-derived repair template was injected along with additional plasmids as previously described^8^ at the following final concentrations: repair template (50ng/μl), Mos1 transposase (pCFJ601, 50ng/μl; under control of Peft-3 promoter), and four markers for negative selection of chromosomal arrays (pMA122 [Phsp-16.41::peel-1], 10ng/μl; pCFJ90 [Pmyo-2::mCherry], 2.5ng/μl; pCFJ104 [Pmyo-3::mCherry], 5ng/μl; and pGH8 [Prab-3::mCherry], 10ng/μl). After injection, worms were allowed to grow for ∼8 days until the plates starved out and no food remained. The starved plates were heat shocked at 34° for 3 hours to kill worms containing extrachromosomal arrays and screened to identify non-fluorescent moving worms as transgene insertion candidates; insertion at the correct locus was confirmed by PCR.

To generate the mScarlet tagged ZEN-4, we adapted a previously published CRISPR RNP-based protocol^28^. A repair template containing mScarlet was made by PCR amplifying the mScarlet sequence using Q5 polymerase (NEB) and primers containing homology arms to the *zen-4* genomic locus on either end with silent mutations to the PAM site to prevent cutting of the repair template by Cas9 (homology arms: upstream 5’-tgttttatttaattcaattcatttcagaaa-3’; downstream 5’-ggacttggtctcgcgatggagttattcctcTCttacgcgacga-3’; silent mutations shown as capitalized letters). PCR amplified mScarlet repair template was purified using a Qiagen Mini-Eluate kit (Qiagen); tracrRNA and crRNAs targeting *zen-4* (5’-AAAATGTCGTCGCGTAAACG-3) and *dpy-10* (co-injection marker; 5’-GCTACCATAGGCACCACGAG-3’) were ordered from IDT and resuspended at a concentration of 200μM. An RNA mix containing duplexes of *zen-4* and *dpy-10* with tracrRNA was made by mixing 4μl *zen-4* crRNA, 1μl *dpy-10* crRNA, and 5μl tracrRNA and incubating at 95°C for 5 minutes followed by 22°C for 5 minutes. An RNP mix was made by mixing 2μl of duplexed RNA mix with 5μl of 40μM Cas9 (Macrolab, Berkeley) and incubating for 5 minutes at room temperature before addition of 0.7μL of 3M KCl (to prevent the Cas9 from precipitating when mixed with repair template). To make the final mix for injection into adult worms, 3μg of repair template was dried via speedvac and resuspended with 3μL of the RNP mix; the mix was spun at maximum speed in a 4°C centrifuge for 15 minutes before injection. Worms from the strain OD3498 were injected, singled and allowed to produce progeny, which were screened for the presence of *dpy-10* co-CRISPR phenotypes (dumpy or roller worms). Dumpy and roller worms were singled and allowed to produce progeny. Isolated dumpy/roller worm populations were genotyped by PCR (oligos: 5’-atccgacctgtgctttgatg-3’, and 5’-TGGACAAAGACGACAAACGAC-3’; wild-type ZEN-4 = 530bp, mScarlet::ZEN-4 = ∼1400bp) to determine for whether mScarlet was inserted into the *zen-4* locus and positive clones were confirmed by visualization of the mScarlet fluorescent marker. Positive clones were confirmed to be homozygous, and *dpy-10* mutations were removed by backcrossing to obtain normally moving non-dumpy worms that produced only normally moving non-dumpy progeny. Strains expressing NMY-2::GFP were generated by using a CRISPR Cas9-based method^29^ to insert a sequence encoding superfolder GFP, which was re-encoded by codon shuffling to render it resistant to *gfp* RNAi targeting the CYK-4::GFP::MEI1N transgene, into the endogenous *nmy-2* locus.

#### Gene knockdown by RNA interference & assessing embryonic viability

RNA interference (RNAi) was performed by injecting dsRNAs targeting the gene of interest into L4 *C. elegans* hermaphrodites, which were subsequently incubated at 16°C or 20°C depending on the experiment. To produce dsRNAs, oligos containing either T3 or T7 promoter sequences were used to amplify the target region of the gene of interest by PCR from either N2 genomic DNA or cDNA. PCR products were purified, either by gel purification or QIAquick PCR Purification kit (Qiagen) and used as the template DNA for 100μl T3 or T7 polymerase reactions (MEGAscript, Invitrogen). Reactions were cleaned using either MEGAclear (Invitrogen) or Qiagen RNeasy (Qiagen) purification kits and eluted in 50μL of nuclease-free water. T3 and T7 reaction products were mixed in equimolar amounts and diluted with 3x soaking buffer (32.7 mM Na_2_HPO_4_, 16.5 mM KH_2_PO_4_, 6.3mM NaCl, 14.1 mM NH_4_Cl) to a final concentration of 1x soaking buffer and annealed (68°C for 10 minutes followed by 37°C for 30 minutes) before measuring the final dsRNA concentration (mg/ml = A_260_ x 0.04) and aliquoting. For targeting a single gene, dsRNA was injected at a concentration of at least 1μg/μl; for double and triple gene depletions, dsRNAs were mixed at equal concentrations to a total concentration of at least 1μg/μl.

For 2-step RNAi in strains expressing a degron-tagged *cyk-4* transgene, L4 hermaphrodites were injected with *cyk-4* dsRNA and incubated at 20°C for 24 hours. These worms were injected a second time with *cyk-4* + *gfp* dsRNAs (mixed at 1:1 concentration) and incubated at 16°C for 18–28 hours before imaging. During imaging of 2-step RNAi embryos, which was done in a room at 20°C, worms were kept on ice with a sheet of bubble wrap in between, thereby maintaining the worms at ∼16°C until immediately prior to dissecting for filming.

To assess brood size, L4 hermaphrodites were injected with dsRNAs and incubated at 20°C for 18 hours, after which they were transferred every 6 hours and the number of embryos laid were counted.

#### Imaging

Embryos for live-imaging experiments were isolated by dissecting gravid adult hermaphrodites in M9 buffer (42 mM Na_2_HPO_4_, 22mM KH_2_PO_4_, 86mM NaCl, and 1mM MgSO_4_), and transferred onto a 2% agarose pad and overlaid with an 18×18mm coverslip. Dissected embryos were imaged using a spinning disk confocal system (either an Andor Revolution XD Confocal System (Andor Technology) with a CSU-10 (Yokogawa) mounted on an inverted microscope (TE200-E; Nikon) with a 60x 1.4NA Plan-Apochromat objective and an EMCCD camera (iXon; Andor Technology), a Zeiss confocal system (Zeiss) with a CSU-X1 (Yokogawa) mounted on an inverted microscope (Axio Observer.Z1; Zeiss) with a 63x 1.4NA Plan Apochromat lens (Zeiss) and a Photometrics QuantEM: 512SC camera, or a Nikon confocal system (Nikon) with a CSU-X1 (Yokogawa) mounted on a Nikon Eclipse Ti2 microscope with a 60x 1.4NA Plan Apochromat lens (Nikon) and an iXon Life EMCCD camera (iXON-L-888; Andor)).

All images were processed, scaled, and analyzed using Fiji ^30^ software.

### QUANTIFICATION AND STATISTICAL ANALYSIS

#### Furrow closure measurements

To image furrow closure over time, an 8 or 9-plane z-series at 2μm intervals was captured every 20s. Imaging was initiated just prior to nuclear envelope breakdown. The largest distance between opposing furrow tips was measured and divided by the initial width starting at anaphase onset to calculate the percent of furrow closure at each timepoint. For montages in the figures, the z-plane in which the furrow was most open throughout the sequence was shown; the z-plane was not adjusted on a per-timepoint basis.

#### Measurement of cortical equatorial mScarlet::ZEN-4

To quantify the cortical recruitment of mScarlet::ZEN-4, worms injected with the specified dsRNAs were dissected 24-37 hours post-injection, depending on the experiment, and 9×2μm z-stacks of the embryos were acquired every 20s with 2×2 binning. The three most coverslip-adjacent cortical z-planes acquired between 260-300s after anaphase onset (defined as the last frame prior to the visible separation of chromosomes) were max projected in time and z to create a single image. Recruitment was quantified by performing a line scan analysis using a 40-pixel long x 30-pixel wide line (∼19.6 x 14.7 µm) centered either on the visible cortical ZEN-4 signal (taking care to ensure the entire line was inside the embryo) or the division plane in the case of embryos without discernable cortical ZEN-4 accumulation. Line scan data was processed by calculating the average background (average signal across the first ∼25% and last ∼25% of the line scan for that embryo) and subtracting this value from the measurements for each point along the line scan. Signal values were normalized by dividing by the mean peak value (average across the central ∼25% of the line scan) across all control embryos. Graphs plot the average normalized cortical mScarlet::ZEN-4 signal ± the 95% confidence interval for each condition.

#### Cortical equatorial mScarlet::ZEN-4 dissolution analysis

For examining the dissolution of centralspindlin cortical localization over time (**Fig. 5B; Fig. S4D**), long image series (extending from anaphase onset through NEBD of the 2^nd^ cell division) were collected of embryos expressing endogenously-tagged mScarlet::ZEN-4 along with untagged WT or GAP Cat^mut^ CYK-4 after RNAi targeting *spd-1*, *nmy-2*, and endogenous *cyk-4*. Sequences were processed and maximum intensity projections of the full z-stacks were generated. Furrowing was evaluated (no furrowing, <50% furrowing, >50% furrowing; numbers of embryos in each class in **Fig. S4A**) and sequences with >50% furrowing were not evaluated further. Cortical localization was compared between embryos expressing WT or GAP Cat^mut^ CYK-4 separately for the pool of embryos that exhibited no furrowing (**Fig. S4D**) and <50% furrowing (**Fig. 5B; Fig. S4D)**. For both the no furrowing and <50% furrowing conditions, movies with weak/no localization, which occurs when the embryo is in the wrong orientation (quantified in **Fig. S4B**) were not analyzed further. Image sequences for all other embryos are presented in the main figure or supplement (**Fig. 5B; Fig. S4D)**.

For the signal quantification shown in Figure 5B, we first calculated the per-pixel background at anaphase onset (the frame immediately preceding visible chromosome segregation, t=0s). Because centralspindlin localizes to the chromosomes and mitotic spindle at this time point, we calculated the per-pixel cytoplasmic background by drawing a 38×50pixel (approximately 18.62×24.5μm) rectangle over the division plane and a smaller 38×25 pixel (18.62×12.25μm) rectangle that encompassed the region containing the chromosomes. A per-pixel background was calculated for the region between the two rectangles [(integrated density in large rectangle-integrated density in small rectangle)/(area of large rectangle – area of small rectangle)]. A corrected fluorescence intensity for each panel was calculated by subtracting the background (per-pixel background * area in pixels of the rectangle) from the integrated intensity in the rectangle (see **Fig. S4C**).

### KEY RESOURCES TABLE

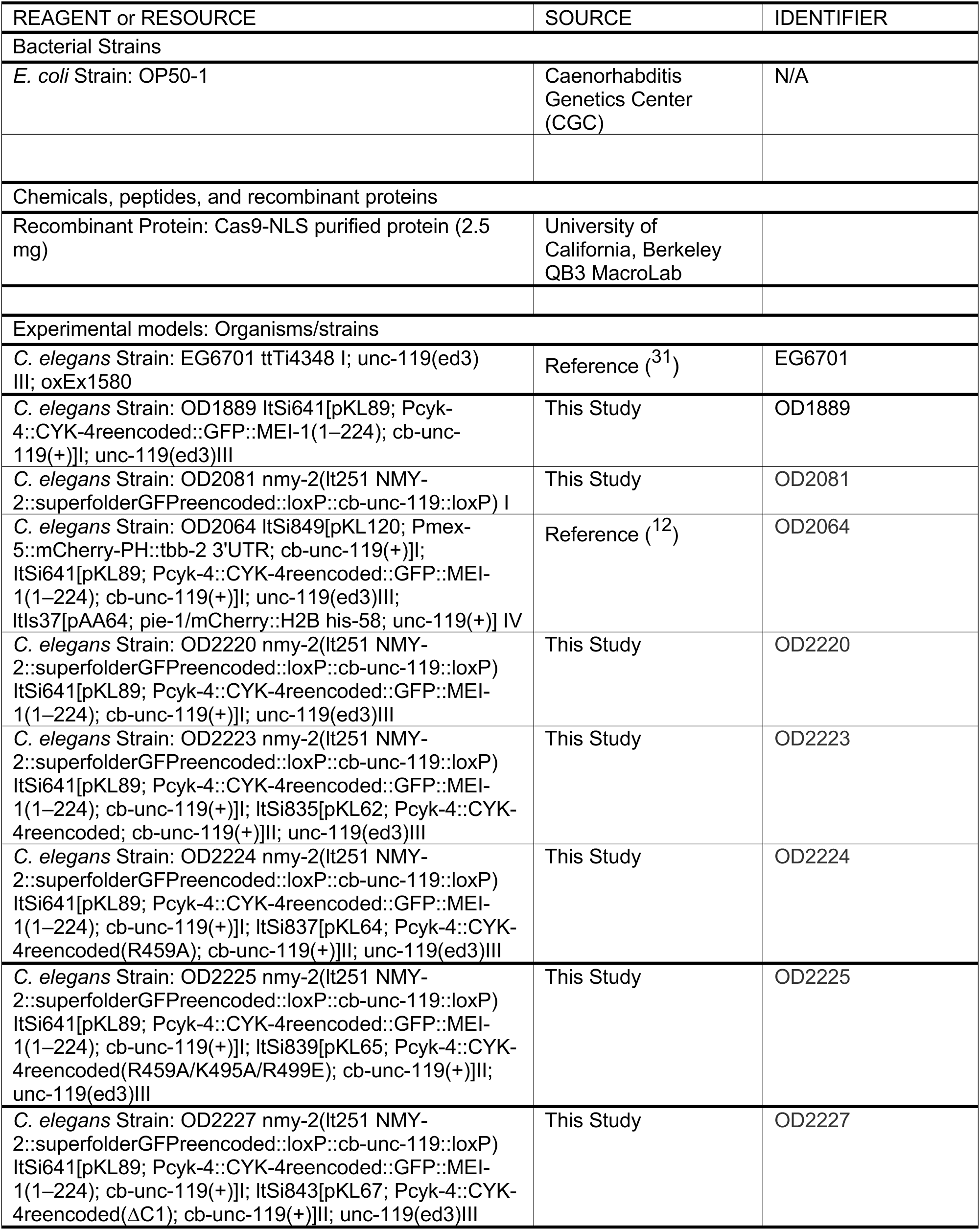
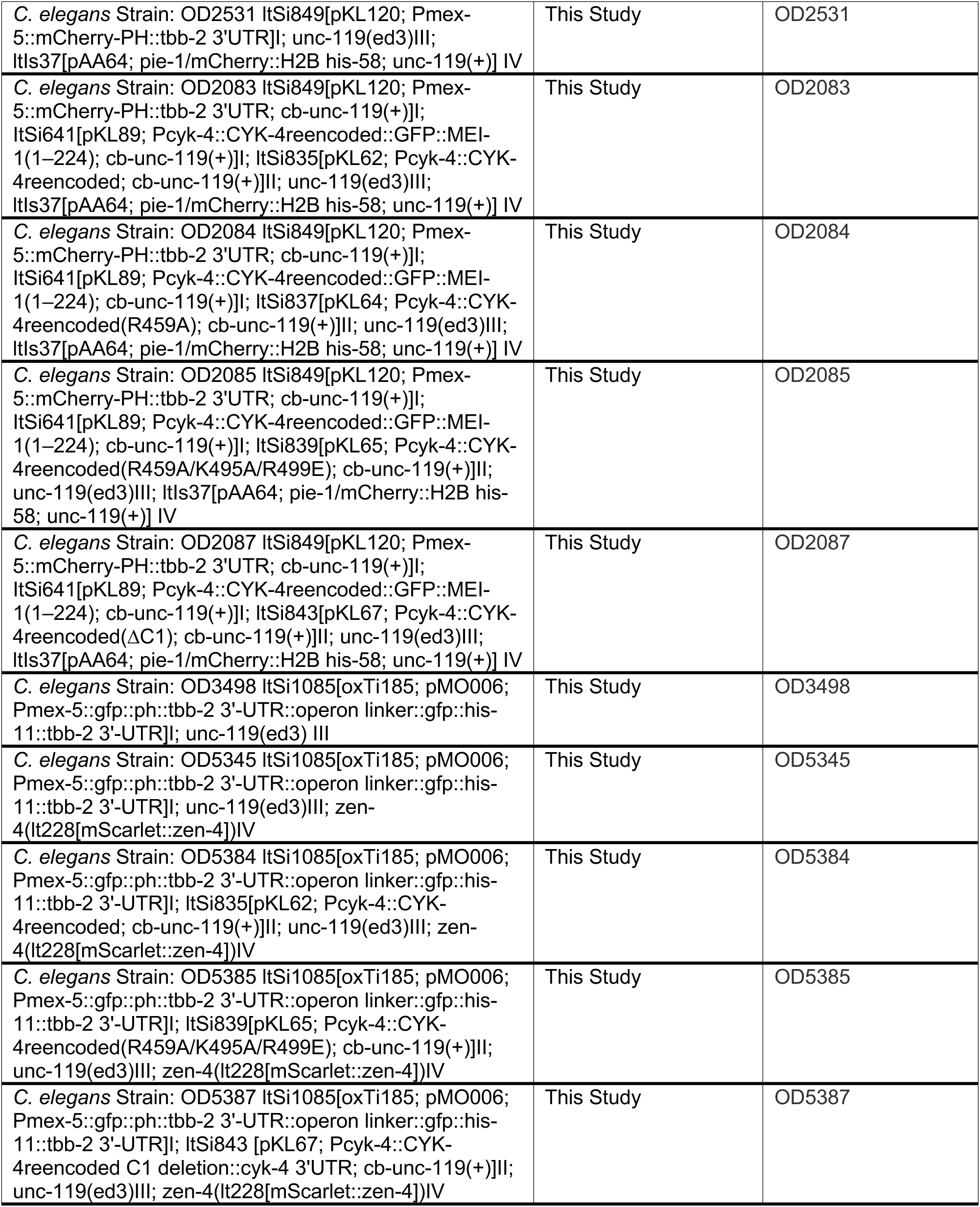
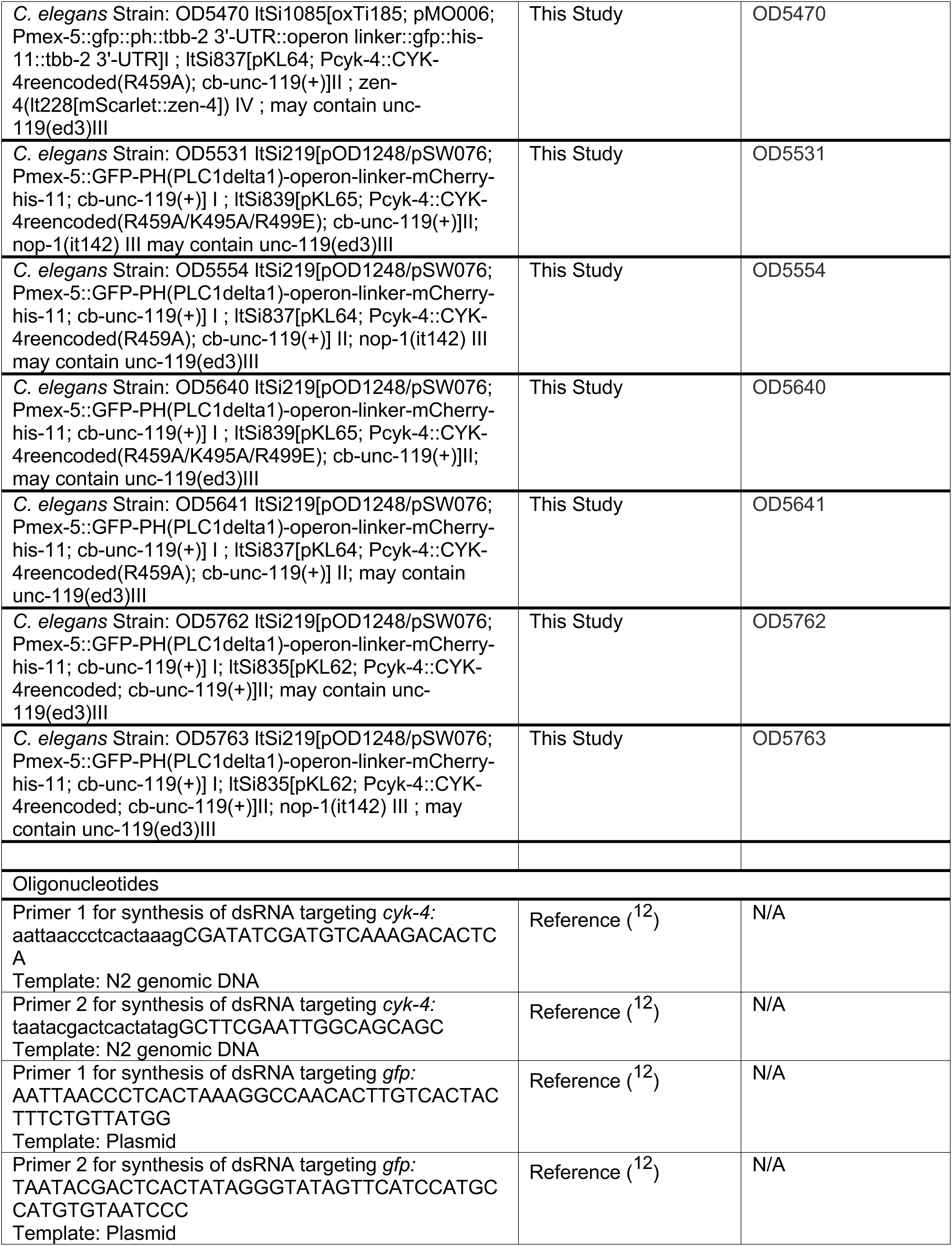
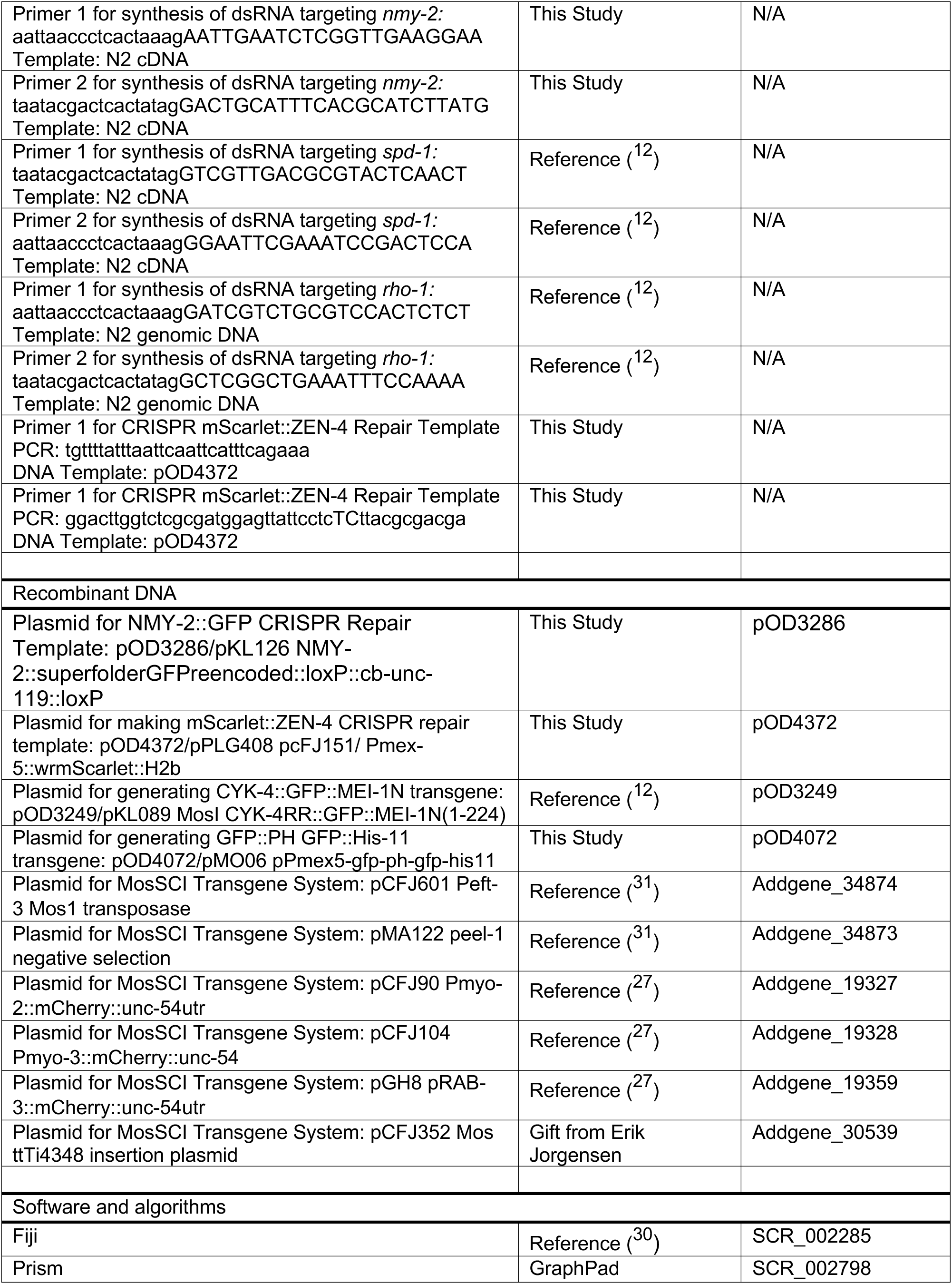
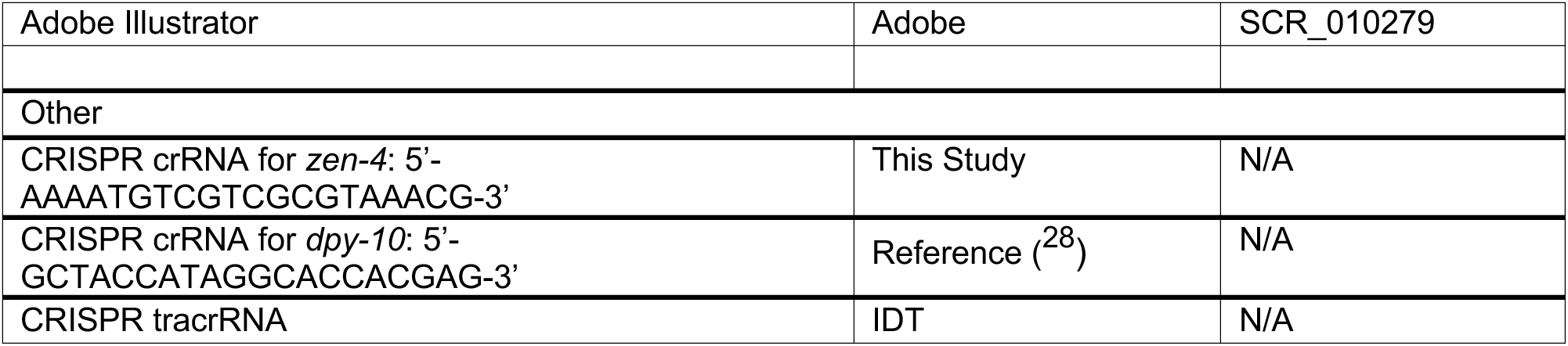

